# Interaction between ciliary component proteins from *Chlamydomonas* revealed by CRISPR/CAS9, cryo-electron tomography and mass spectrometry

**DOI:** 10.1101/2024.04.02.587733

**Authors:** Leo Luo, Noemi Zimmermann, Akira Noga, Alexander Leitner, Takashi Ishikawa

## Abstract

To understand molecular mechanism of ciliary beating motion, knowledge of location, interaction and dynamics of >400 component proteins are indispensable. While recent progress of structural biology revealed conformation and localization of >100 proteins, we still need to investigate their networking, art of their interaction and assembly mechanism. We applied CRISPR/CAS9 genome editing technique to the green algae *Chlamydomonas* to engineer a deletion mutant of a ciliary component, FAP263, located at the distal protrusion, and examined it structurally by cryo-electron tomography (cryo-ET) and mass spectrometry (MS). Cryo-ET and atomic model fitting demonstrated that the FAP263 deletion mutant lacks additional components, FAP78, and FAP184. Unassigned density near FAP263 in the cryo-ET map of WT cilia is likely FAP151, as suggested by cross-linking mass spectrometry. Based on the structure, we modeled how these four proteins might form a complex. Furthermore, it was shown that dynein f phosphorylation is inhibited in the FAP263 mutant, indicating an important role of this protein complex for dynein f phosphorylation. Our study demonstrates a novel approach to investigate protein networking inside cilia.

## Introduction

Motile cilia, beating organelle to cause either swimming of the cells or extracellular fluid flow, are composed of more than 400 proteins (Pazour *et al*, 2005). Ciliary motion is considered to be product of dynamic interactions and networking of these proteins. There has been wide variety of attempts to reveal the protein interactions, 3D arrangements and their dynamic changes. Recent progress of single particle cryo-EM analysis and folding prediction by Alphafold2 allowed visualization of >100 components which exist in the peripheral microtubule doublet with 96nm periodicity (Walton *et al*, 2023), components of the central pair apparatus with 32nm periodicity (Gui *et al*, 2022), and intraflagellar transport complexes (Hesketh *et al*, 2022). Cryo-electron tomography of intact cilia, combined with high resolution obtained by single particle cryo-EM, enabled modeling of conformational change of dynein and associated proteins during motion (Zimmermann *et al*, 2023). Further expansion of our knowledge to cover more component proteins at various states of beating motion and at various stages of ciliogenesis is awaited both for basic understanding of ciliary movement and medical investigation of ciliopathy (Wallmeier *et al*, 2020).

Both structural and functional studies utilize deletion mutants. By comparing the wild type and deletion mutants, or by comparing the wild type and a strain with a deletion mutant rescued by a tagged gene, location of the target protein can be detected based on decrease or increase of density in the 3D map, respectively. In cilia research, combination of deletion mutants and cryo-ET revealed functional roles and locations of a number of component proteins in intact cilia (Ishikawa, 2016). In the case target proteins are small and hard to detect as a loss of density in deletion mutants or in case deletion could cause collapse of other proteins in the complex, genetic tag to increase density helps efficiently. Components of the radial spoke (Oda *et al*, 2014c), the dynein regulatory complex (Oda *et al*, 2014b), ruler proteins for the doublet microtubule (Oda *et al*, 2014a) and an inner dynein scaffold protein (Kutomi *et al*, 2021), were located by cryo-ET in this way.

However, availability of deletion mutants for motile cilia research was severely limited. *Chlamydomonas reinhardtii* has been the most popular model organism for motile cilia research because of abundance of mutants of ciliary components, which were isolated based on motility defect. Active mutagenesis of *Chlamydomonas* by targeting gene of interest, however, is complicated because homologous recombination of this species is not established. While there are wide variety of deletion mutants, first induced chemically or by radiation and later by random insertion (Li *et al*, 2016), systematic methods to delete targeted gene have been unavailable for years. Meanwhile *Tetrahymena thermophila*, another popular model organism for motile cilia research can be mutated in targeting genes using homologous recombination, which has been used for biochemical research by purifying mutated proteins (Ichikawa *et al*, 2015). For cellular studies such as cellular cryo-ET of cilia, mutation of *Tetrahymena* cannot be used easily, since there are 50 copies of each gene in the small nucleus and complete exclusion of the intrinsic sequence is not straightforward.

In this study, we generated deletion mutant of a ciliary component FAP263 from *Chlamydomonas*, which is located near the outer surface of the A-tubule and dynein f, using CRISPR/CAS9. We characterized these deletion mutants biochemically and structurally. Our study demonstrates proof-of-principle of combination of CRISPR/CAS9 and cryo-ET for cilia research, as well as its advantage to study influence of gene deletion to other component proteins.

## Results and discussion

We made deletion mutant of FAP263 by inserting stop codon to *Chlamydomonas* genome by CRISPR/CAS9 following the protocol of (Shin *et al*, 2016). Mutation was confirmed by PCR and sequencing (Fig. 1).

**Fig. 1.**
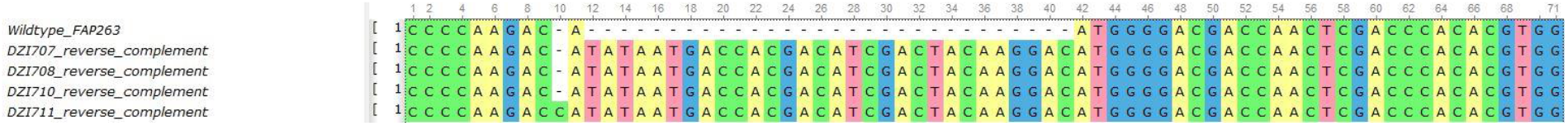
Sequence of the FAP263 deletion mutant by CRISPR/CAS9.

After back-crossing the mutant to WT to minimize a risk of off-target effects, we further structurally analyzed FAP263 deletion mutant by cryo-ET and subtomogram averaging (Figs.2, 3A, Supplementary Fig. 1). Protein components can be located in the cryo-ET map by fitting high-resolution single particle cryo-EM structure of split doublet microtubule from *Chlamydomonas* cilia (Walton *et al*, 2023). There is an area, where density exists in WT, but not in the FAP263 deletion mutant (Fig. 2; Supplementary Fig. 1) which likely corresponds to proteins lost by FAP263 deletion. In the fitted atomic model from single particle cryo-EM, this lost area corresponds to FAP78/FAP184 and FAP263 (Fig. 3B). Therefore this density can be explained as a complex of FAP263, FAP78 and FAP184.

**Fig. 2.**
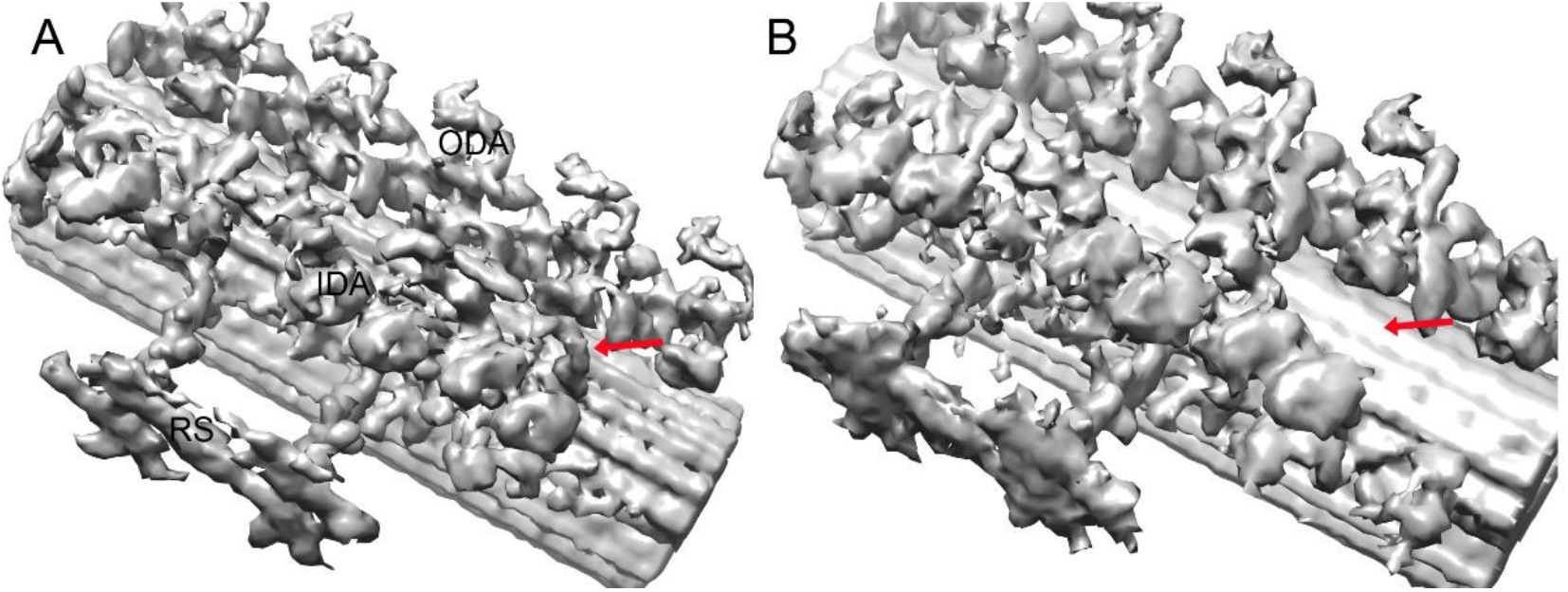
Cryo-ET structure of FAP263 deletion mutant. The maps by subtomogram averaging from cryo-ET of WT (A) and the FAP263 deletion mutant (B). The distal protrusion (in WT) and the corresponding place (in the deletion mutant) are indicated by red arrows. Other major components (ODA: outer dynein arms; IDA: inner dynein arms; RS: radial spoke) are also indicated.

**Fig. 3.**
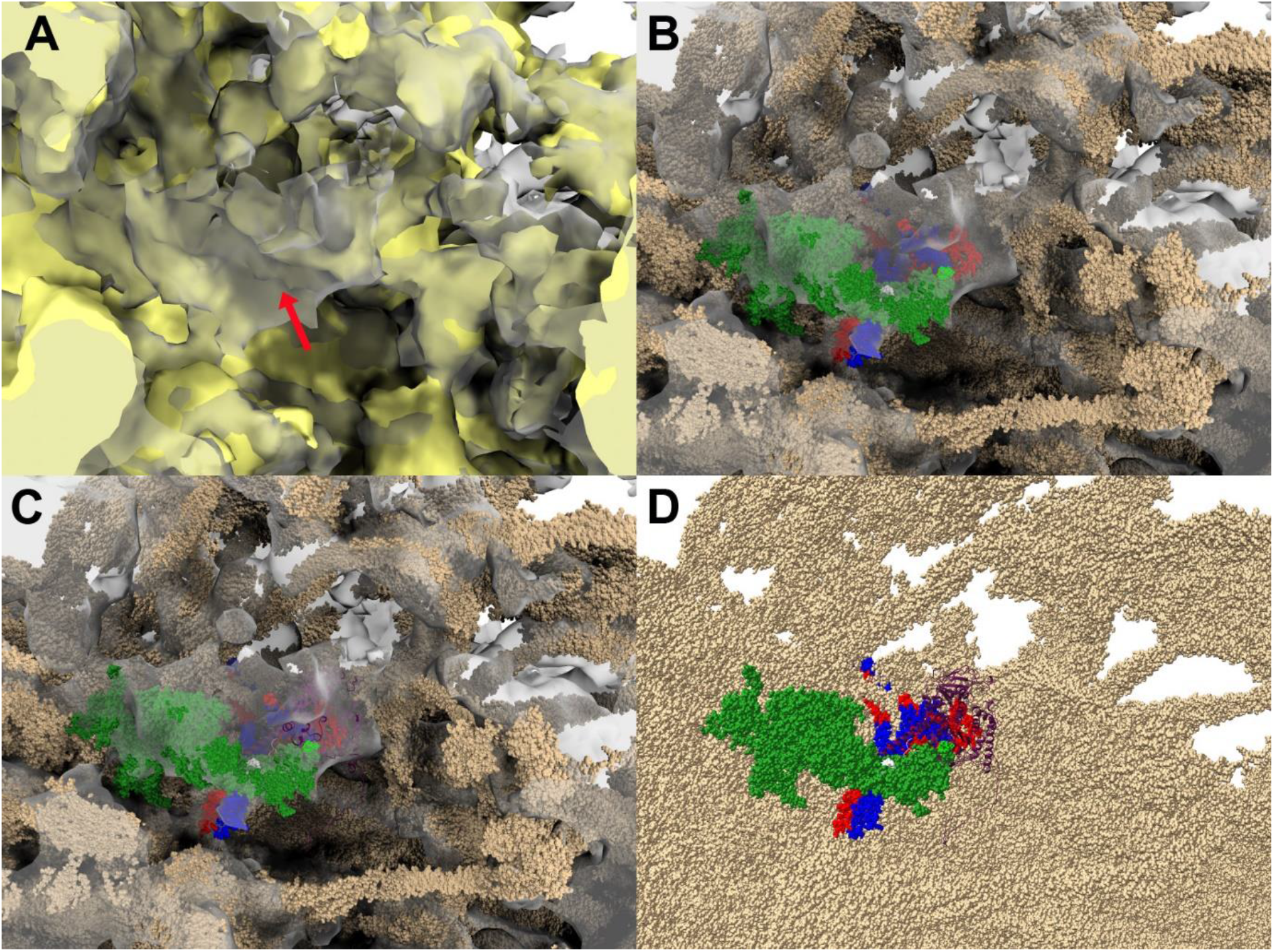
Enlarged view of the distal protrusion including FAP263 and associated proteins in cilia from averaged cryo-ET map of WT (grey) and the FAP263 deletion mutant (yellow). In (A) the density difference, corresponding to the distal protrusion, is indicated by a red arrow. In (B), atomic models from (Walton *et al*, 2023) are fitted to the cryo-ET map and superimposed. FAP78 (green), FAP184 (blue), FAP283 (red). (C) FAP151, modeled by Alphafold2 is added in purple. (D) Only atomic models are presented from (C). The color code is the same as (C).

However we found a small unassigned region as well (Fig. 3B). This unassigned density is positioned between FAP78 and FAP263. To find candidates for an additional protein located in this region, we performed cross-linking mass spectrometry (Leitner *et al*, 2014) and looked for proteins that were identified in earlier proteomics work on *Chlamydomonas* cilia (Pazour *et al*, 2005), but were not localized in the cryo-EM structure. The cross-link data points to a putative interaction between FAP78 and the protein FAP151 (Fig. 4), which was not involved in the list of single particle cryo-EM analysis. It should be pointed out that the confidence of this identification is relatively low, which may be attributed to the low abundance of the protein complex in cilia, and/or specific mass spectrometric properties of the cross-linkned peptides. Nevertheless, we modeled FAP151 by Alphafold2 and fitted to the unassigned density (Fig. 3B). FAP151 fits well to other components and is likely a piece of this complex (Fig. 3CD). Since FAP78 is solved in full-length by single particle cryo-EM (Walton *et al*, 2023), this density may contain a part of FAP78 as well.

**Fig. 4.**
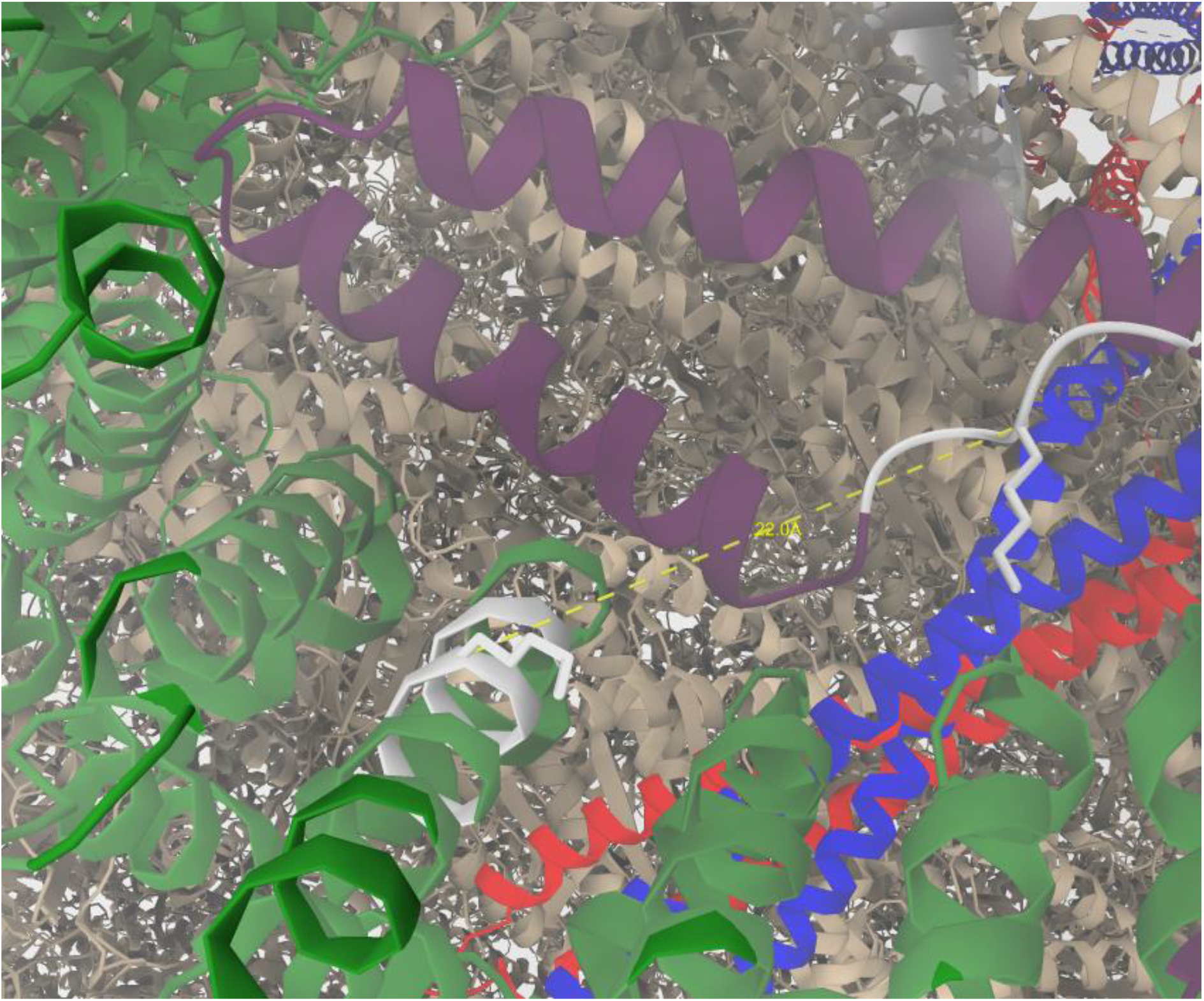
Crosslinking MS sites, 366K of FAP78 (green) and 355K of FAP151 (purple), are shown in white.

Furthermore, we performed phosphoproteomics experiments to study the role of protein phosphorylation on putative functions of the subcomplex involving FAP263. Comparative mass spectrometric analysis of phosphopeptides from wild type and FAP263 mutant samples after enrichment with titanium dioxide (Leitner *et al*, 2010) resulted in a high coverage of the *Chlamydomonas* phosphoproteome. Four proteins in the distal protrusion have more than one assigned spectrum for peptides that are detected phosphorylated in WT but not in the FAP263 mutant (Table 2). Among dynein isoforms, dynein f (DHC10) is located close to FAP263 and likely influenced by its deletion for phosphorylation. Indeed dynein f shows phosphorylation in WT, but not upon FAP263 deletion (Table 1). This suggests that the FAP78/FAP151/FAP184/FAP263 complex is responsible for dynein f phosphorylation. Among these four proteins, FAP78 is likely to have kinase activity, according to classification by Panther (https://phytozome-next.jgi.doe.gov/report/gene/Creinhardtii_v5_6/Cre12.g536600). However, phosphorylation by another protein located near this complex, for example a Nima kinase CNK4, cannot be excluded.

**Table 1.** Phosphorylation of dyneins.

| Gene symbol / uniprot accession number | Average number of peptide spectrum matches for phosphopeptides in control | Average number of peptide spectrum matches for phosphopeptides in FAP263 mutant |
| --- | --- | --- |
| DHC1 A0A2K3D1N8 | 0 | 0.3 |
| DHC2 A0A2K3DE97 | 2.7 | 2.3 |
| DHC3 A0A2K3DN16 | 23.7 | 28 |
| DHC4 A0A2K3E2Q0 | 21.7 | 31.3 |
| DHC5 A0A2K3E2P7 | 4.7 | 0.6 |
| DHC6 A0A2K3DSC5 | 2.3 | 0 |
| DHC7 A0A2K3CYE9 | 6.3 | 9.7 |
| DHC8 A0A2K3CUR8 | 11.7 | 14 |
| DHC9 A0A2K3E486 | 2 | 2.3 |
| DHC10 A0A2K3CY97 | 1.3 | 0 |
| DHC11 A0A2K3D5V4 | 13 | 19.3 |
| DHC12 A0A2K3DQQ4 | 25.7 | 43.3 |
| DHC13 A0A2K3DV97 | 41.3 | 31 |
| DHC14 A8J1M5 | 1.3 | 0.3 |
| DHC15 A0A2K3D8D3 | 0 | 0 |
| DHC16 A0A2K3DLV6 | 0 | 0 |

**Table 2.** Phosphorylation of possible distal protrusion proteins.

| Gene symbol / uniprot<br>accession number | Average number of identified<br>phosphopeptides in control | Average number of identified<br>phosphopeptides in FAP263<br>mutant |
| --- | --- | --- |
| FAP263 A0A2K3CQ22 | 5.7 | 0 |
| FAP184 A0A2K3DWC0 | 48.7 | 2.7 |
| FAP78 A0A2K3D574 | 11.3 | 2.7 |
| FAP151 A0A2K3D802 | 1 | 0 |
| CNK4 Q6UPR2 | 1.7 | 1 |

Although we have not seen visual difference of swimming property between WT and the FAP263 deletion mutant, change of beating frequency and amplitude caused by mutation of Ccdc113/Ccdc96 (corresponding to FAP263/184) in *Tetrahymena* was reported (Bazan *et al*, 2021). While phosphorylation of IC138, associated with dynein f, was reported (Bower *et al*, 2009), phosphorylation of dynein f itself has not been studied. Functional roles of this complex is still to be investigated.

In this study, we demonstrated CRISPR/CAS9 of *Chlamydomonas* is useful for structural research of cilia. The FAP263 deletion causes loss of FAP78, FAP151, and FAP184. This approach can be applied to solve various unanswered questions, such as precise location of various species of dyneins, role of individual dyneins and regulatory proteins in ciliary motion, as well as ciliogenesis.

## Materials and methods

### Strains and media

*C. reinhardtii* strain CC-503 cw92 mt+ (Chlamydomonas Resource Center, https://www.chlamycollection.org/) were used in this study. Cells were grown in Tris-acetate-phosphate (TAP) medium in a constant light/dark cycle (light cycle: 21:00-11:00, dark cycle 11:00-21:00).

### crRNA design

crRNAs were chosen using Benchling (Hirano *et al*, 2019). On-target and off-target scores calculated by Benchling against *Chlamydomonas r*. genome were aimed to be above 60 and 90, respectively. crRNAs were purchased from Integrated DNA Technology’s (IDT) online custom Alt-R® CRISPR-Cas9 guide RNA tool.

### Design of repair template and preparation

The repair template was designed to have two homology arms upstream and downstream of the cutting side, each with a length of 25 bp. In the middle of the homology arms is a FLAG-tag with two stop codons. The FLAG-tag serves the detection of mutant colonies by polymerase chain reaction (PCR). The repair template was ordered as a forward and reverse oligonucleotide on Microsynth AG. The oligonucleotides were ordered with 3 phosphorothioate bonds on both 5’ and 3’. To anneal the oligonucleotides, 1 μL of forward and 1 μL of reverse oligonucleotides were mixed in 18 μL IDT RNA duplex buffer. Afterwards the mix was heated to 95 °C for 2 minutes and cooled down at 0.1 °C/sec to room temperature.

### Paromomycin cassette for selection

The plasmid pSI103-1, which confers paromomycin resistance, was ordered from Chlamydomonas Resource Center. The resistance cassette was amplified using chemically competent *E. coli*. The plasmid was extracted with a plasmid extraction kit from QIAGEN. Afterwards, the DNA is linearized by KpnI-HF from Biolabs. The linearized DNA was concentrated to 1 μg/μL by ethanol precipitation.

### Preparation of Cas9/gRNA RNP

gRNA was assembled by mixing 5 μL of 100 μM IDT Alt-R® crRNA with 5 μL of 100 μM IDT Alt-R® tracrRNA. The mix was incubated at 95 °C for 2 minutes and cooled down at 0.1 °C/sec to room temperature. 4.5 μL of 10 μM gRNA was mixed with 4.5 μL of 10 μM IDT Alt-R® Cas9, 1.5 μL of 10x NEB 3.1 and 4.5 μL of ddH_2_O. The RNP system was incubated at 37 °C for 15 minutes.

### Transformation by electroporation

150 mL of cw92 cells were grown for 3 days in TAP medium. Cells were centrifuged at 800 g for 5 minutes. The cells were washed once with electroporation buffer (30 mM Hepes, 5 mM MgSO_4_, 50 mM K-acetate, 1 mM Ca-acetate, 60 mM Sucrose). Afterwards the cells were pelleted again and resuspended in 2-3 times the volume of cells in electroporation buffer. Cells were diluted to a concentration of 3 * 10^8^ cells/mL. Then they were incubated at 40 °C for 30 minutes at 120 rpm. 100 μL of heat shocked cells, 15 μL of RNP, 4.5 μL of repair template and 1 μg of pSI103-1 were mixed in a cuvette from Bio-Rad (Catalog No. 165-2086). The mixture was electroporated using ECM® 630 from BTX – Harvard Apparatus with the following conditions: 410 V low voltage, Resistor 25 Ohm, Capacitance 600 uF. After electroporation cells were incubated for 1 hour at 15 °C. The cells were transferred into 10 mL of 60 mM TAP sucrose for recovery overnight under constant light and shaking.

### Transfer on TAP agar plates

After recovery we centrifuged the cells at 800 g for 5 minutes and plated them on 1.5% TAP agar plates with 10 μg/mL paromomycin. Colonies can be picked after 5-7 days and transferred to 96 well plate with TAP. Confirmation of mutant colonies was done by PCR using Phusion™ High-Fidelity DNA Polymerase. Screening for mutants were done by gel electrophoresis. To confirm mutants, they were sent for sequencing at Microsynth AG.

### Cell culture and harvesting cilia

Chlamydomonas cell culture and cilia isolation were followed by Witman’s protocol (Witman, 1986).

### Cryo-ET

Cryo-ET grid preparation and data acquisition were after our previous work (Zimmermann *et al*, 2023), using one-sided manual blotting and freezing by Cryo-plunge (Gatan, USA) and the Titan Krios G3 transmission electron microscope (TFS, USA) with Quantum energy filter (Gatan). Subtomogram average using pseudo nine-fold symmetry and 96nm periodicity of cilia was carried on using our algorithm published previously (Bui & Ishikawa, 2013; Zimmermann *et al*, 2023).

### Cross-linking mass spectrometry

*C. reinhardtii* strain cc124-was cultured for 3 days. Cilia were isolated by dibucaine. Isolated cilia by dibucaine were treated with 1% OGP in equal volume to remove cell membrane. cOmplete™ Proteasehemmer-Cocktail by Roche were used to stop protein degradation. Protein concentration was measured using BCA assay and adjusted to 0.5-2 mg/ml with a total amount of 50-100 μg protein. Cross-linking experiments were performed at room temperature with the amine-reactive disuccinimidyl suberate in isotopically light and heavy form for 1 hour (DSS-d_0_/d_12_, Creative Molecules). Afterwards the protocol by (Leitner et al., 2014) was used to process the samples further. Samples were analyzed by liquid chromatography-tandem mass spectrometry on an Orbitrap Fusion Lumos instrument (ThermoFisher Scientific), and MS data was analyzed by xQuest.

### Phosphoproteomics

*C. reinhardtii* strain cc124-was used as wild type control for this experiment. Three replicates of the cc124-strain and three replicates of the FAP263 mutant strain were cultured in 300 ml of TAP medium. Each culture was maintained for a duration of three days. Isolation of cilia and sample preparation were done as described above. To prevent dephosphorylation, PhosSTOP™ from Sigma Aldrich was added to sample. Amount of protein was adjusted to 150 μg per sample by BCA assay. Mass spectrometric analysis of protein phosphorylation was performed following (Leitner et al., 2010) using Titansphere TiO material (GL Sciences) for enrichment. Samples were analyzed by liquid chromatography-tandem mass spectrometry on an Orbitrap Fusion Lumos instrument (ThermoFisher Scientific), and MS data was analyzed by FragPipe/MSFragger. For data analysis a cutoff of Peptide Prophet Probability > 0.95 was chosen. For comparison of phosphorylation states between wild type and FAP263 mutant the total amount of peptide spectrum matches of phospho peptides were counted.

## Author contribution

LL conducted experiments, including CRISPR/CAS9, cryo-ET of the FAP263 deletion mutant, MS. NZ did cryo-ET of WT. NA backcrossed mutant cells to WT. AL supervised and designed MS. TI designed the entire project. The manuscript was prepared by LL and TI.

## Acknowledgement

We appreciate Prof. Kenichi Wakabayashi for advices at start-up of CRISPR/CAS9, ScopeM and CEMK (ETHZ) for cryo-EM support and Prof. Paula Picotti (ETH Zurich) for access to the laboratory infrastructure and instrumentation. This research was funded by grants from Swiss National Science Foundation (IZLIZ3_200294, 310030_192644), NanoArgovia (FunkEM project) and Novartis Biomedical Foundation (to TI).

## Figure caption

**Supplementary Fig. 1.**
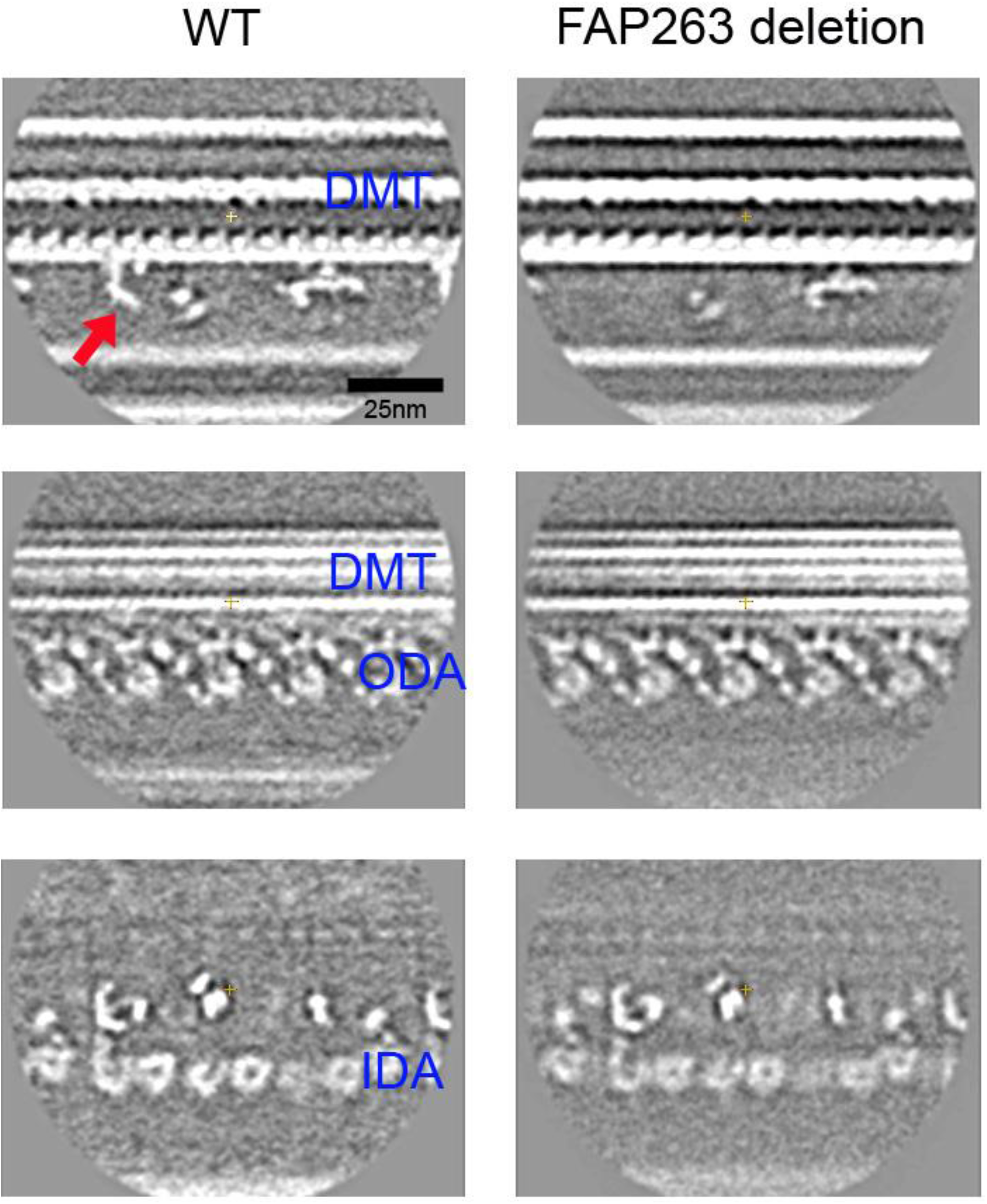
Cross sections from averaged subtomograms from cryo-ET of WT (left) and FAP263 deletion mutant (right). In the top row, presence and absence of the distal protrusion is indicated by red arrows. In the middle and bottom rows, the parallel sections including doublet microtubules (DMT) shows outer dynein arms (ODA) and inner dynein arms (IDA) in the same structure between WT and the mutant.

