## Supplementary Fig. 1 for "Interaction between ciliary component proteins from *Chlamydomonas* revealed by CRISPR/CAS9, cryo-electron tomography and mass spectrometry"

Supplementary Fig.1 Cross sections from averaged subtomograms from cryo-ET of WT (left) and FAP263 deletion mutant (right). In the top row, presence and absence of the distal protrusion is indicated by red arrows. In the middle and bottom rows, the parallel sections including doublet microtubules (DMT) shows outer dynein arms (ODA) and inner dynein arms (IDA) in the same structure between WT and the mutant.

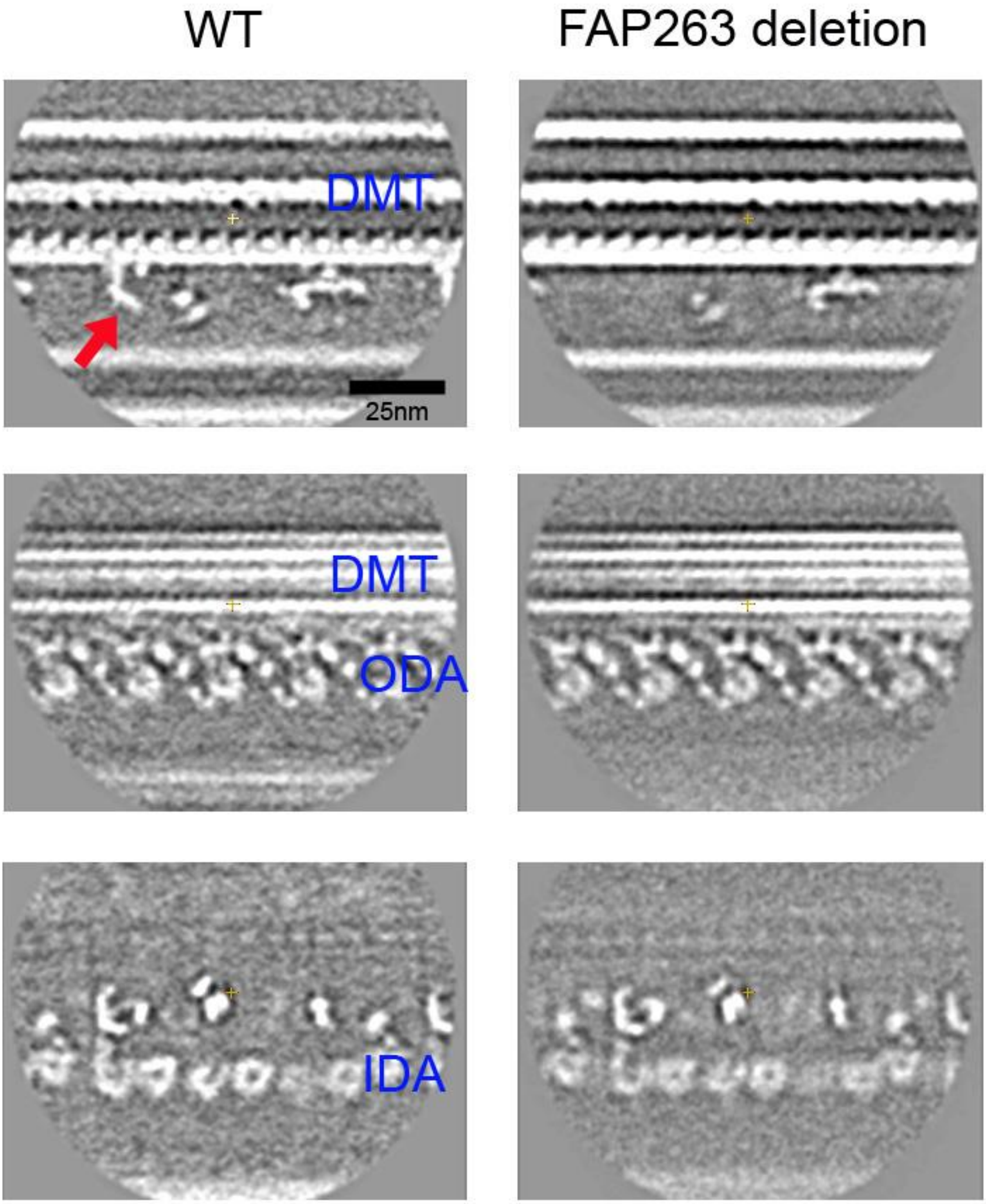
